# Effects of age on pre-stimulus neural activity predictive of successful memory encoding: an fMRI study

**DOI:** 10.1101/2020.05.18.102939

**Authors:** E. Song Liu, Joshua D. Koen, Michael D. Rugg

## Abstract

Pre-stimulus subsequent memory effects (SMEs) – differences in neural activity preceding the onset of study items that are predictive of later memory performance – have consistently been reported in young adults. The present fMRI experiment investigated potential age-related differences in pre-stimulus SMEs. During study, healthy young and older participants made one of two semantic judgments on images, with the judgment signaled by a preceding cue. In the test phase, participants first made an item recognition judgment and, for each item judged old, a source memory judgment. Age-invariant pre-stimulus SMEs were observed in left dorsomedial prefrontal cortex, left hippocampus, and right subgenual cortex. In each case, the effects reflected lower BOLD signal for later recognized items, regardless of source accuracy, than unrecognized items. A similar age-invariant pattern was observed in left orbitofrontal cortex, but the effect in this region was specific to items attracting a correct source response compared to unrecognized items. In contrast, the left angular gyrus and fusiform cortex demonstrated negative pre-stimulus SMEs that were exclusive to young participants. The findings indicate that age differences in pre-stimulus SMEs are regionally specific and suggest that pre-stimulus SMEs reflect multiple cognitive processes, only some of which are vulnerable to advancing age.

---

Episodic memory - memory for unique personal events - declines disproportionately with advancing age relative to other forms of memory (Nilsson, 2003; Nyberg et al, 2012). A key factor responsible for age-related episodic memory decline is a reduction in the efficacy of processes that support successful memory encoding (Craik & Byrd, 1982; Werkle-Bergner et al, 2006; Old & Naveh-Benjamin, 2008; Craik & Rose, 2012; Friedman & Johnson, 2014). Accordingly, numerous studies have used functional magnetic resonance imaging (fMRI) to identify neural correlates of age-related declines in the efficacy of memory encoding. The great majority of such studies have employed the “subsequent memory procedure” (Wagner, 1998; Brewer et al, 1998; Paller & Wagner, 2002), in which neural activity elicited by study items is contrasted according to the judgment given the items on a subsequent memory test. These studies have consistently demonstrated that, in young adults, neural activity during encoding differs according to later memory performance, phenomena referred to as subsequent memory effects (SMEs). The effects of age on SMEs have been well-studied over the past 20 years or so (e.g. Morcom et al., 2003; Gutchess et al., 2005; Dennis et al., 2008; Miller et al., 2008; de Chastelaine et al., 2011, 2016; Mormino et al., 2012; Park et al., 2013; Mattson et al., 2014; for review see Maillet & Rajah, 2014). Age differences have been reported to be robust in regions typically showing so-called ‘negative’ SMEs (regions demonstrating reduced BOLD signal for subsequently remembered relative to forgotten items), whereas age differences are more modest and identified less consistently in regions demonstrating ‘positive’ SMEs (Maillet & Rajah, 2014).

Of importance, there is robust evidence that neural activity during the seconds *prior to* the presentation of a study item is also predictive of subsequent memory performance (e.g. Otten et al., 2006; 2010; Adcock et al., 2006; Park & Rugg, 2009; Addante et al., 2015; de Chastelaine and Rugg, 2015; Cohen at al., 2019; for review see Cohen et al., 2015). The existence of pre-stimulus SMEs suggests that a full understanding of memory encoding, and how it differs with age, will require elucidation not only of the neural processing engaged during the online experience of an event, but also of the processing that precedes it.

fMRI studies of pre-stimulus SMEs have mainly employed paradigms in which study items are preceded by a cue that provides information about the upcoming study event, such as the format or modality of the study item (e.g. Addante et al., 2015; Park & Rugg, 2009) or the nature of the upcoming study judgment (e.g., de Chastelaine & Rugg, 2015). In these paradigms, pre-stimulus SMEs are examined by contrasting the neural activity occupying the cue-item interval according to the judgment accorded the item on the subsequent memory test. The studies have typically reported greater pre-stimulus BOLD signals in the hippocampus and surrounding medial temporal lobe (MTL) preceding study items that went on to be recollected (recognition was accompanied by retrieval of qualitative information about the study event) relative to items that were either recognized on the basis of an acontextual sense of familiarity, or that were unrecognized. Similar ‘positive’ pre-stimulus SMEs have also been reported in several cortical regions, including dorsolateral prefrontal, lateral parietal and posterior midline cortex (Addante et al., 2015).

Although the majority of fMRI studies have described exclusively positive pre-stimulus SMEs, there is evidence that the direction of the effects can be influenced by task demands (de Chastelaine & Rugg, 2015). de Chastelaine & Rugg (2015) reported a reversal in the direction of pre-stimulus SMEs as a function of the nature of the upcoming encoding task, identifying positive hippocampal pre-stimulus SMEs for study items subjected to a semantic (animacy) judgment and negative pre-stimulus SMEs for items subjected to an orthographic (syllabic) judgment. Although there is no correspondence between the directionality of fMRI and event-related potential (ERP) effects, it is noteworthy that Koen et al. (2018) reported a condition-dependent reversal of ERP pre-stimulus SMEs (for similar findings, see Padovani et al., 2011). Specifically, for briefly presented (300 ms) studied items, Koen and colleagues identified typical negative-going pre-stimulus SMEs (cf. Otten et al., 2006). When study items were presented for 1000 ms, however, positive-going pre-stimulus SMEs were observed.

Pre-stimulus SMEs have been interpreted in two principal ways. One interpretation is that they reflect controlled processes elicited by the pre-stimulus cue that support the adoption of a specific task set or the engagement of task-appropriate preparatory processes (for review, Cohen et al, 2015). According to such ‘preparatory’ accounts (e.g., Park & Rugg, 2009; de Chastelaine & Rugg, 2015; Addante et al, 2015; Koen et al, 2018), more effective preparation leads to more efficient processing of the study item, freeing up cognitive resources that can then be devoted to encoding operations. Alternately, pre-stimulus SMEs might reflect spontaneous fluctuations in neural states that are more or less conducive to effective encoding (e.g., Fernandez et al, 1999; Otten et al., 2002; Yoo et al., 2012; Salari & Rose, 2016; Schneider & Rose, 2016; Sweeney-Reed et al., 2016; Ezzyat et al., 2017; Cohen et al, 2019; Sadeh et al, 2019). Notably, in an uncued experimental design, Yoo et al (2012) reported that later remembered study items (scenes) were associated with lower levels of pre-stimulus fMRI BOLD activity in the parahippocampal cortex than later forgotten scenes. By employing ‘real-time’ fMRI analyses (see Salari & Rose, 2016 for a similar analysis applied to EEG), these researchers went on to demonstrate that subsequent memory was enhanced when study scenes were presented during periods in which the parahippocampal cortex was relatively quiescent, a finding that could be taken to imply that negative pre-SMEs reflect neural states conducive to successful encoding. These two accounts of the functional significance of pre-stimulus SMEs are, of course, not mutually exclusive.

Research on age differences in pre-stimulus SMEs is currently sparse, restricted to studies employing electrophysiological measures, and has led to mixed findings. Koen and colleagues (2018) reported that ERP pre-stimulus SMEs were markedly more prominent in young than older adults. This finding was interpreted as evidence that the older adults did not engage in effective preparation for the upcoming study event, an account consistent with a broader body of research suggesting that older adults do not spontaneously engage ‘proactive’ control. By contrast, an EEG study by Strunk & Duarte (2019) reported age-invariant effects in time-frequency measures of pre-stimulus neural activity elicited by pre-stimulus cues that signaled the modality of the upcoming study item. Contrary to the conclusions of Koen et al. (2018), these authors proposed that older and young adults were equally capable of engaging pre-stimulus anticipatory processes that facilitate episodic encoding.

In light of these sparse and seemingly contradictory findings, the present fMRI study further addressed the question of the effects of age on pre-stimulus SMEs. We employed an experimental procedure very similar to one first described by Mattson and colleagues (2014), modifying the design to permit pre-and post-stimulus BOLD signals to be deconvolved (cf. Park & Rugg, 2009; de Chastelaine & Rugg, 2015). The procedure was adopted because it permits assessment of the neural correlates of the encoding of information supporting subsequent source and item memory judgments while controlling for the potentially confounding effects of memory strength (Mattson et al. 2014; see also Rugg et al, 2012). In light of prior findings (e.g. Adcock et al., 2009; Park & Rugg, 2009; de Chastelaine & Rugg, 2015), we expected to identify pre-stimulus SMEs in young participants in the MTL and, possibly, in cortical regions also (Addante et al., 2015). At issue is whether analogous findings are evident in older individuals. To the extent that pre-stimulus SMEs identified with fMRI reflect the engagement of preparatory (proactive) processes - processes that, as noted above, have been held to decline with advancing age (Braver et al, 2009; Braver, 2012) - we expected to observe an age-related attenuation of pre-stimulus SMEs.

## Materials and Methods

This study was conducted with the approval of the Institutional Review Boards of the University of Texas Southwestern Medical Center and the University of Texas at Dallas. All participants provided written consent prior to each experimental session. They were compensated at the rate of $30 per hour and refunded for travel expenses.

### Participants

A total of 55 healthy adults, comprising 28 young adults (14 females) aged between 18 and 30 years (mean age: 23 years) and 27 older adults aged between 65 and 77 years (mean age: 69 years) contributed to the data analyzed below. Participants were recruited from the University of Texas at Dallas and the greater Dallas metropolitan area. All participants were right-handed, with normal or corrected-to-normal vision. Individuals were excluded if they reported a history neurological, psychiatric, cardiovascular disease, or any contraindication for MRI (for additional exclusion criteria based on cognitive performance, see *Neuropsychological testing* below). Neuroimaging data from an additional 5 participants were collected but excluded from analyses due to an insufficient number of trials in critical experimental conditions (2 young and 1 older adult), a programming error (1 older adult), and an incidental structural MRI finding (1 older adult).

### Neuropsychological testing

All participants undertook a battery of standardized neuropsychological tests, which was administered on a separate day prior to the MRI session. The battery (see Table 1) included the Mini-Mental State Examination (MMSE), the California Verbal Learning Test-II (CVLT; Delis et al., 2000), Wechsler Logical Memory Tests 1 and 2 (Wechsler, 2009), Trail Making tests A and B (Reitan and Wolfson, 1985), the Symbol Digit Modalities test (SDMT; Smith, 1973), the F-A-S subtest of the Neurosensory Center Comprehensive Evaluation for Aphasia (Spreen and Benton, 1977), the WAIS–R subtests of forward and backward digit span (Wechsler, 1981), a category fluency test (Benton, 1968), Raven’s Progressive Matrices (List 1, Raven et al., 2000) and the Wechsler Test of Adult Reading (WTAR; Wechsler, 1981). To reduce the likelihood of including participants with mild cognitive impairment, potential participants were excluded if they scored < 27 on the MMSE, > 1.5 SD below age norms on any standardized memory test, > 1.5 SD below age norms on two or more standardized non-memory tests, or if their estimated full-scale IQ was < 100.

**Table 1.** Demographic data and neuropsychological test results (means and SDs) for young and older adults.

|  | *Young Adults* | *Older Adults* | *p* |
| --- | --- | --- | --- |
| *N (male/female)* | 14/14 | 13/14 | - |
| *Age (years)* | 22.36 (2.78) | 68.74 (2.89) | - |
| *Education (years)* | 15.25 (1.95) | 16.59 (2.52) | 0.032 |
| *MMSE* | 29.11 (0.92) | 28.96 (0.90) | 0.558 |
| *CVLT short delay free recall* | 13.07 (2.51) | 10.44 (3.31) | 0.002 |
| *CVLT short delay cued recall* | 13.25 (2.47) | 11.67 (2.65) | 0.026 |
| *CVLT long delay free recall* | 13.57 (2.66) | 11.33 (2.75) | 0.003 |
| *CVLT long delay cued recall* | 13.86 (2.43) | 11.96 (2.43) | 0.006 |
| *CVLT recognition hits* | 15.36 (1.13) | 14.70 (1.51) | 0.077 |
| *CVLT recognition false alarms* | 1.04 (1.90) | 2.26 (2.23) | 0.033 |
| *WMS logical memory I* | 31.68 (8.22) | 27.89 (4.29) | 0.037 |
| *WMS logical memory II* | 28.75 (7.46) | 25.93 (4.93) | 0.103 |
| *Forward/backward digit span* | 18.89 (4.49) | 18.67 (3.93) | 0.843 |
| *Digit-symbol substitution test* | 62.43 (10.40) | 47.48 (6.37) | 0.001 |
| *Trail Making test A (sec)* | 21.66 (6.75) | 29.46 (9.52) | 0.001 |
| *Trail making test B (sec)* | 48.92 (21.00) | 66.96 (23.50) | 0.004 |
| *FAS fluency* | 44.82 (11.37) | 46.11 (9.98) | 0.656 |
| *Category fluency* | 24.25 (4.20) | 21.37 (4.08) | 0.013 |
| *WTAR (raw)* | 39.56 (5.08) | 40.81 (5.02) | 0.361 |
| *RAVEN's List I* | 11.18 (0.82) | 9.37 (1.69) | 0.001 |

### Experimental Materials

Experimental stimuli were presented using Psychophysics Toolbox 3.0 (http://psychtoolbox.org) implemented in Matlab 2017b (www.mathworks.com). Participants viewed centrally presented stimuli over a gray background via a mirror mounted above the scanner head-coil. The critical experimental stimuli comprised 270 images of every-day objects, food items, and animals. Stimuli were selected randomly and without replacement to create 28 different 180-item study lists that were administered to yoked pairs of young and older participants. Within each study list, the images were randomly separated into 5 blocks (36 items per block). Critical stimuli for the test phase included all of the images from the study phase, along with an additional 90 randomly selected ‘new’ images (giving a total 270 images). Study and test lists were pseudorandomized such that participants were not presented with more than three consecutive trials containing the same image class. Prior to the study phase, participants received practice on 32 images (8 for practice 1, 8 for practice 2, and 16 for practice 3, see below) outside the scanner. Prior to the test phase, practice was provided using the 16 images from practice 3 intermixed with 8 new images.

### Procedure

#### Study Phase

The scanned study phase is schematized in Figure 1. Each trial began with the presentation of a centrally located green fixation cross for 500 ms. This was followed by a pre-stimulus cue: either a red “X” or “O”, which was presented at fixation for a duration of 750 ms. The cue was followed by a white fixation cross, which was presented for either 1500 ms, 3500 ms, or 5500 ms, (rectangular distribution). The study image onset concurrently with the termination of the white cross and was presented for 1500 ms. The image was followed by a second white fixation cross which again varied randomly in duration for 1500 ms, 3500 ms, or 5500 ms. Participants were instructed to make their judgment about the study image (see below) as rapidly but as accurately as possible and before the termination of the fixation cross.

**Figure 1.**
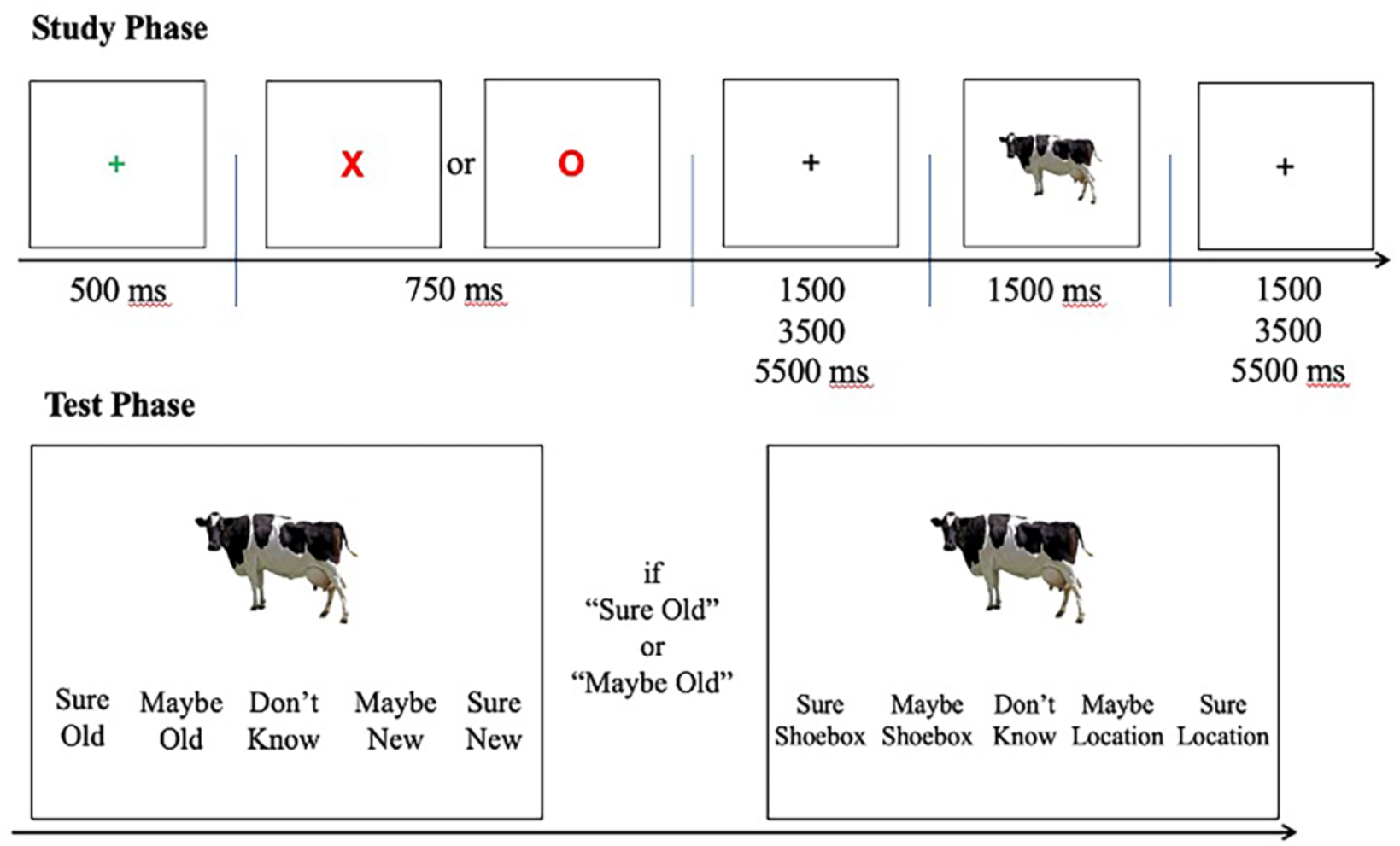
Schematic of the encoding task and subsequent memory test. At encoding, participants were instructed to indicate whether the item depicted by the image “fits into a shoebox” (X – shoebox trials); or “is more likely to be found indoors or outdoors” (O – location trials).

The pre-stimulus cues signaled the judgment to be made on the upcoming image. An “X” signaled the requirement to make a ‘shoebox’ judgment (“does the item depicted by the image fit into a shoebox?”), and an “O” signaled that a ‘location’ judgment should be made (“is the item depicted by the image more likely to be found indoors or outdoors?”). Participants were instructed to use different hands (counterbalanced across participants) to make the two judgments. The index and middle fingers were associated with “Indoors/Fit inside a shoebox” and “Outdoors/Does not fit inside a shoebox”, respectively.

Before entering the scanner, participants were given detailed instructions about the study tasks and completed 3 practice study phases. The first practice was self-paced with feedback. This was followed by a timed practice block with feedback, followed by a practice block that was identical to the actual study phase (and hence did not include feedback). Feedback was restricted to whether the correct hand was employed for each task judgment and did not include judgment accuracy.

#### Test Phase

Participants were administered a surprise memory test outside of the scanner approximately 15 minutes after the completion of the study phase (hence, the study phase engaged incidental rather than intentional encoding processes). They were instructed to make memory judgments accompanied by a confidence rating (see Figure 1). The test task first required a five-way item memory judgment (“Sure Old”, “Maybe Old”, “Don’t know”, “Maybe New”, “Sure New”). For each item endorsed “Old” participants went on to make a source judgment concerning the study task associated with the item using the response alternatives “Sure Location”, “Maybe Location”, “Don’t Know”, “Maybe Shoebox”, and “Sure Shoebox”. For both judgments, participants were encouraged to use the entire range of confidence ratings. Both test judgments were required within a 10 sec window following item onset, after which a trial timed out and the following trial was initiated. Three rests breaks, comprising intervals of 1 min, 5 min, and 1 min respectively, were provided between each quadrant of the test phase.

#### Behavioral data analysis

Trials that received multiple responses, no response, or a response with the incorrect hand during the study phase were excluded from behavioral and subsequent fMRI analyses. We computed four behavioral dependent measures that were subjected to group analyses: median study reaction time (RT), item recognition accuracy (Pr), response bias (Br), and source memory accuracy (pSR). Median study RTs were computed for the three memory bins used for the fMRI analysis (see below), and these RTs were analyzed with a 2 (age group) X 3 (memory) mixed design ANOVA.

The remaining three measures (Pr, Br, and pSR) were computed from the memory test performed outside of the scanner. Following Mattson et al. (2014), item memory accuracy (Pr) was computed as the difference between hit rate of study items regardless of source memory accuracy and false alarm rate to new items. Item response bias was estimated using the *Br* metric (*pFA / [1 − (pHit -pFA)]*; Snodgrass & Corwin, 1988). Overall source accuracy (pSR) was derived from a guessing-corrected single high threshold model (Snodgrass & Corwin, 1988; Park & Rugg, 2009; Mattson et al, 2014) using the formula:

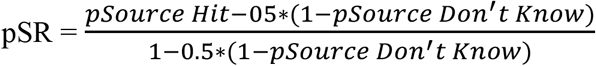

Age differences in the pR, pSR, and Br measures was investigated using independent samples *t*-tests that did not assume equal variance between the young and older adult groups.

### MRI Data Acquisition

Functional and anatomical images were acquired from a 3T Philips Achieva MRI scanner (Philips Medical Systems) equipped with a 32 channel receiver head coil. Functional images were acquired with a T2* weighted blood oxygen level dependent (BOLD) multiband echoplanar (EPI) sequence (flip angle = 70°, FOV = 200 * 240 mm, TR = 1.5 ms, TE = 30 ms, multiband factor = 2). EPI volumes comprised 44 slices (inter-slice gap of 0.5 mm) with isotropic 2.5 mm voxels. Slices were acquired in an interleaved order and oriented parallel to the AC-PC line. Anatomical images were acquired with a T1-weighted magnetization-prepared rapid gradient echo (MPRAGE) pulse sequence (FOV = 256 × 256 mm, 1×1×1mm isotropic voxels, 176 slices, sagittal acquisition).

### fMRI Data Analysis

#### fMRI data preprocessing

The functional data were preprocessed with Statistical Parametric Mapping (SPM12, Wellcome Department of Cognitive Neurology, London, UK) implemented in Matlab 2017b (www.mathworks.com). The functional data were reoriented, subjected to a two-pass realignment procedure with images realigned to the first image of a session and then re-aligned to the mean EPI image, and corrected for slice acquisition time differences by using *sinc* interpolation with reference to the time of acquisition. Finally, images were spatially normalized to a study-specific EPI template following previous published procedures (de Chastelaine et al., 2011; 2016), and smoothed with an 8 mm full-width at half-maximum kernel. The data from the five study sessions were concatenated using the *spm_concatenate.m* function and prior to the first level general linear model (GLM).

#### fMRI analyses

The functional data were analyzed with a two-stage mixed effects model. In the first level GLM, data for each participant were modeled to capture cue-and stimulus-related neural activity. Cue-related activity was modeled by convolving a canonical hemodynamic response function with a variable boxcar function that extended from cue onset until the onset of the study item with (Park & Rugg, 2009). Stimulus-elicited neural activity was modeled by convolving a canonical hemodynamic response function with a delta function time-locked to the onset of the study item. Following Mattson et al. (2014), the design matrix included three events of interests for both cue-and stimulus-elicited activity: 1) source hits – studied items that later attracted correct and highly confident item and source memory judgments; 2) source misses – studied items that later attracted correct and highly confident item memory judgments, but incorrect, low confidence or “don’t know” source judgments, and 3) item misses – studied items that later attracted a new, low confidence old, or “Don’t Know” item memory response. The inclusion of low confidence old responses (for which accuracy was barely above chance) in the item miss bin was to ensure a sufficient number of trials for fMRI analyses. Source hit and source miss trials were restricted to studied items that later received high confidence item judgments so as to mitigate the confounding of item memory strength with source memory accuracy (cf. Squire et al., 2007; Rugg et al., 2012). The first level GLM also included regressors to model events of no interest (studied items later attracting correct but low confidence item judgments, and trials where no response or multiple responses were given) in a single regressor, six motion regressors, spike covariates for volumes showing transient displacement > 1 mm or 1° in any direction, and constants modelling mean signal in each scan session.

To identify voxels that were sensitive to the different memory conditions in an unbiased manner, participant-specific parameter estimates for cue-elicited activity were taken forward to a second level analysis and subjected to a 2 (age group) by 3 (memory condition) mixed-design ANOVA model. The regions of interest (ROIs) employed in subsequent analyses were defined as clusters identified by the main effect of memory condition in the aforementioned ANOVA (the ANOVA did not identify any clusters that demonstrated an age x memory condition interaction at the whole brain thresholds specified below). The ANOVA had a height threshold of p <.005 and clusters were deemed significant if they exceeded the p<.05 FWE corrected cluster extent threshold (k > 90) determined by SPM’s false discovery rate (FDR) correction for multiple comparisons (Chumbley et al., 2010). For each significant cluster, mean parameter estimates were extracted from 5mm radius spheres centered on each of the top three peak voxels (separated by a minimum of 7 mm). The parameter estimates were averaged across the peaks and subjected to 2 (age group) by 3 (memory condition) ANOVAs. Effects were deemed significant at p<.05 after Geisser-Greenhouse correction for non-sphericity. Note that whereas a main effect of memory condition is a foregone conclusion in these ANOVAs given the criteria for ROI selection, the patterning of the effects between the levels of the memory factor and interactions between memory and age group are free to vary and are not biased by the ROI selection procedure.

Because of our *a priori* interest in and predictions for the hippocampus (see Introduction), data from this region were subjected to small volume rather than whole brain correction (voxel-wise FWE corrected height threshold of p<.05). The hippocampus was traced on the mean sample-specific anatomical scan and used to create a bilateral mask that defined the volume within which correction was applied (see de Chastelaine & Rugg, 2015, for a similar approach).

#### Time course estimation

To visualize pre-stimulus effects, time-courses of the BOLD response in each of the regions identified by the foregoing analyses were estimated using a finite impulse response (FIR) model (cf. Park & Rugg, 2009; de Chastelaine & Rugg, 2015). The time-courses were estimated across 10 time points (sampling interval of 1.5s) that began 7.5 seconds prior to the onset of the study item and extended 6 seconds post-stimulus. FIRs were estimated for each memory condition of interest (source hit, source miss, and item miss) and collapsed across the three cue-stimulus intervals (2.25, 4.25, and 6.25 s).

## Results

### Neuropsychological Test Performance

Table 1 reports demographic information and neuropsychological test scores for the two age groups. Older adults performed significantly worse than young adults on tests of declarative memory, reasoning, processing speed and category fluency. The patterning of the neuropsychological test scores across the age groups is consistent with that reported previously for similar populations (e.g., Park et al, 2001; de Chastelaine et al., 2011). The greater educational experience of the older group reflects the fact that the majority of the young participants were undergraduate college students whereas the majority of the older sample had obtained college degrees.

### Behavioral Performance

#### Study Phase

##### Study RT

Table 2 reports median study RTs as a function of age group and the subsequent memory conditions employed in the fMRI analyses. A 2 (age group) by 3 (memory condition) ANOVA revealed no significant interaction between age group and memory condition (F_1.97, 104.54_ = 1.09, p = 0.34, partial-η^2^ = 0.02), and nor was there any significant main effect of memory condition (F_1.97, 104.54_ = 0.99, p = 0.37, partial-η^2^ = 0.02). However, there was a main effect of age group (F_1, 53_ = 16.88, p < 0.001, partial-η^2^ = 0.24), reflecting faster responses on the part of the young participants.

**Table 2.** Mean and SD for study RT (in ms) segregated by subsequent memory condition and age group

|  | *Young adults* | *Older adults* |
| --- | --- | --- |
| *Source hit* | 1036 (173) | 1269 (248) |
| *Source miss* | 1041 (238) | 1291 (298) |
| *Item miss* | 1000 (180) | 1284 (295) |

#### Test Phase

##### Item Memory

Item memory performance is summarized in Table 3. Item recognition accuracy (Pr) did not differ significantly across the age groups (young: M = 0.70, SD = 0.13; older: M = 0.69, SD = 0.13; t_53_ = 0.13, p = 0.89, Cohen’s d = 0.04). This was also the case when Pr was estimated for highly confident judgments only (young: M = 0.64, SD = 0.16; M = older: 0.68, SD = 0.14; t_53_ = 0.92, p = 0.36, Cohen’s d = 0.27).

**Table 3.** Mean and SD for the proportions of item memory judgments for old and new trials by age group and confidence rating.

|  | *Young Adults* |  | *Older Adults* |  |
| --- | --- | --- | --- | --- |
|  | *Old items* | *New items* | *Old items* | *New items* |
| *Confident Old* | 0.68 (0.16) | 0.02 (0.05) | 0.75 (0.13) | 0.07 (0.06) |
| *Unconfident Old* | 0.11 (0.10) | 0.07 (0.09) | 0.04 (0.06) | 0.06 (0.07) |
| *Don't Know* | 0.05 (0.06) | 0.07 (0.10) | 0.04 (0.04) | 0.06 (0.06) |
| *Unconfident New* | 0.07 (0.05) | 0.27 (0.22) | 0.04 (0.04) | 0.12 (0.15) |
| *Confident New* | 0.10 (0.10) | 0.58 (0.31) | 0.14 (0.09) | 0.73 (0.22) |

Turning to response bias (Br), there was no significant difference between the age groups in bias for item memory judgments collapsed across confidence (young: M = 0.27, SD = 0.24; older: M = 0.29, SD = 0.17; t_53_ = 0.28, p = 0.78, Cohen’s d = 0.08). However, Br for highly confident judgments was significantly more liberal (although still conservative) in the older group (young: M = 0.11, SD = 0.15; older: M = 0.22, SD = 0.15; t_53_ = 2.62, p = 0.01, Cohen’s d = 0.71).

##### Source Memory

Source memory performance is summarized in Table 4. In light of the substantial prior evidence of age-related decline in source memory (for review, Spencer & Raz, 1995; Old & Naveh-Benjamin, 2008; Koen & Yonelinas, 2014), age differences in source memory were evaluated with a one-tailed (directional) t-test. This revealed that source accuracy (pSR) was significantly higher in the young age group (young: M = 0.58, SD = 0.17; older: M = 0.51, SD = 0.14; t_53_ = 1.77, p = 0.04, Cohen’s d = 0.48).

**Table 4.**
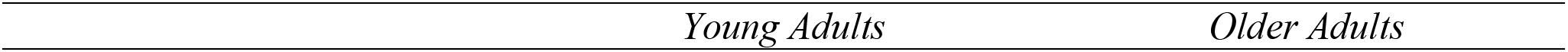
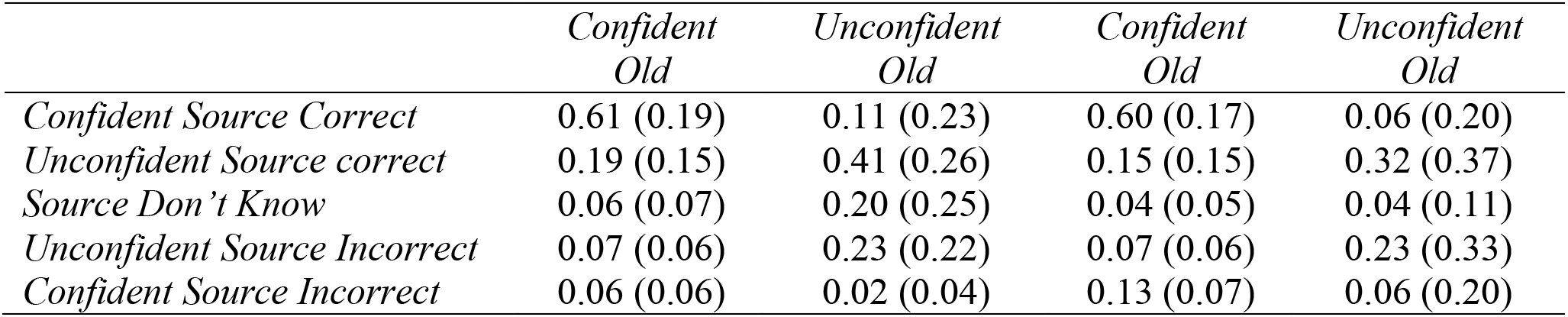
Means and SDs for the proportion of source memory judgments for correctly recognized study items.

| *Young Adults* | *Older Adults* |
| --- | --- |

|  | *Confident*<br>*Old* | *Unconfident*<br>*Old* | *Confident*<br>*Old* | *Unconfident*<br>*Old* |
| --- | --- | --- | --- | --- |
| *Confident Source Correct* | 0.61 (0.19) | 0.11 (0.23) | 0.60 (0.17) | 0.06 (0.20) |
| *Unconfident Source correct* | 0.19 (0.15) | 0.41 (0.26) | 0.15 (0.15) | 0.32 (0.37) |
| *Source Don't Know* | 0.06 (0.07) | 0.20 (0.25) | 0.04 (0.05) | 0.04 (0.11) |
| *Unconfident Source Incorrect* | 0.07 (0.06) | 0.23 (0.22) | 0.07 (0.06) | 0.23 (0.33) |
| *Confident Source Incorrect* | 0.06 (0.06) | 0.02 (0.04) | 0.13 (0.07) | 0.06 (0.20) |

Following Mattson et al (2014), we also contrasted the proportions of confident correct source judgments made to confidently recognized items. These proportions did not significantly differ between the age groups (t_53_ = 1.81, p = 0.08, Cohen’s d = 0.49; see Table 4). Of importance, older adults were however significantly more likely to make confident *incorrect* source judgments for items confidently endorsed as old (see Table 4; t_53_ = 4.16, p < 0.001, Cohen’s d = 1.13).

### fMRI results

#### Events of Interest

The mean (SD; range) trial numbers for source hits, source misses and item misses were, respectively, 75 (35; 10-135), 22 (12; 8 – 58), and 55 (28; 15-115) in the young group, and 80 (29; 21-135), 32 (12; 13-115) and 44 (23; 8-64) in the older group. While there was no significant age difference in the trial counts for source hits and item misses (minimum *p* = .10), the trial count for source misses was significantly higher in the older group (t_53_ = 2.92, p = 0.01, Cohen’s d = 0.79).

#### Whole-Brain Analysis

As detailed in Tables 5 and 6 and illustrated in Figure 2, whole brain analysis revealed significant main effects of subsequent memory in five cortical regions: left dorsomedial PFC (dmPFC), orbitofrontal cortex, angular gyrus, fusiform gyrus, and right subgenual cortex (as already noted, the whole brain age by memory condition interaction contrast did not identify any clusters that survived FWE correction). The mean values of the parameter estimates derived from each region (see Methods) were subjected to 2 (age group) by 3 (memory condition) ANOVAs (see Methods). As noted previously, while a main effect of memory condition is a foregone conclusion in each of these ANOVAs, the patterning of the effects across source hit, source miss, and item miss trials and their moderation by age group are free to vary. The results of the ANOVAs revealed non-significant interactions between age and memory condition in left dmPFC and orbitofrontal cortex, and in right subgenual cortex (see Table 6 and Figure 2), indicative of age-invariant pre-stimulus SMEs in those regions. Pairwise contrasts of the data collapsed across age groups revealed that the pre-stimulus SMEs in left dmPFC and right subgenual cortex took the form of lower activity for source hits and source misses relative to item misses. By contrast, pre-stimulus activity in left orbitofrontal cortex demonstrated a more graded pattern, with lower activity for source hits than for item misses, but with neither source hits nor item misses differing significantly from source misses. As evident in Figure 2, there were outlying data points for both the left orbitofrontal and right subgenual cortex (> 2.5 SD above or below the across-group mean). Removal of these outliers did not alter the findings reported above.

**Table 5.** Regions showing a main effect of memory condition in the whole-brain ANOVA. For each region, MNI co-ordinates of the main peak (bolded) and two sub-peaks are listed.

| Region | MNI co-ordinates |  |  | cluster size | peak z |
| --- | --- | --- | --- | --- | --- |
|  | x | y | z |  |  |
| *Left Dorsomedial PFC (dmPFC)* | **-13** | **58** | **23** | 946 | 4.41 |
|  | -13 | 46 | 25 |  |  |
|  | -3 | 36 | 58 |  |  |
| *Left Orbitofrontal Cortex* | **-23** | **33** | **-15** | 158 | 3.74 |
|  | -28 | 31 | -8 |  |  |
|  | -41 | 21 | -15 |  |  |
| *Right Subgenual Cortex* | **12** | **26** | **-10** | 98 | 3.60 |
|  | -6 | 21 | -20 |  |  |
|  | -6 | 13 | -10 |  |  |
| *Left Angular Gyrus* | **-38** | **-85** | **35** | 91 | 3.83 |
|  | -38 | -80 | 30 |  |  |
|  | -46 | -77 | 40 |  |  |
| *Left Fusiform Gyrus* | **-58** | **-47** | **-18** | 92 | 4.68 |
|  | -58 | -32 | -23 |  |  |
|  | -61 | -35 | -18 |  |  |

**Table 6.** ANOVA interaction contrasts and follow-up pairwise t tests on parameter estimates extracted from clusters demonstrating a main effect of memory condition in the whole brain analysis.

| Region | Age x memory interaction |  |  | Pairwise t tests |  |  |
| --- | --- | --- | --- | --- | --- | --- |
| | F | p | Partial $\eta^2$ | Source Hit vs Source Miss | Source Miss vs Item Miss | Source Hit vs Item Miss |
| *L. Dorsomedial PFC (dmPFC)* | $F_{(1.92, 101.65)} = 0.01$ | 0.99 | < 0.01 | *n.s.* | $t = -4.41$,<br>$p < 0.001$ | $t = -2.88$,<br>$p = 0.001$ |
| *L. Orbitofrontal* | $F_{(1.61, 85.59)} = 3.01$ | 0.07 | 0.05 | *n.s.* | *n.s.* | $t = -5.13$,<br>$p < 0.001$ |
| *R. Subgenual* | $F_{(2.00, 105.90)} = 0.07$ | 0.93 | < 0.01 | *n.s.* | $t = -4.54$,<br>$p < 0.001$ | $t = -3.60$,<br>$p < 0.001$ |
| *L. Angular Gyrus* | $F_{(1.69, 89.64)} = 4.45$ | **0.02** | 0.08 | Young: *n.s.*<br>Old: *n.s.* | Young: $t = -3.75$,<br>$p < 0.001$<br>Old: *n.s.* | Young: $t = -4.94$,<br>$P < 0.001$;<br>Old: *n.s.* |
| *L. Fusiform Gyrus* | $F_{(1.92, 101.76)} = 6.76$ | **< 0.001** | 0.11 | Young: *n.s.*<br>Old: *n.s.* | Young: $t = -4.57$,<br>$p < 0.001$<br>Old: *n.s.* | Young: $t = -4.85$,<br>$p < 0.001$;<br>Old: *n.s.* |

*Note:* Significance levels of the pairwise t tests were corrected for multiple comparison within each region (3 contrasts) using the Holm-Bonferroni method. The corrected p-values are reported.

**Figure 2.**
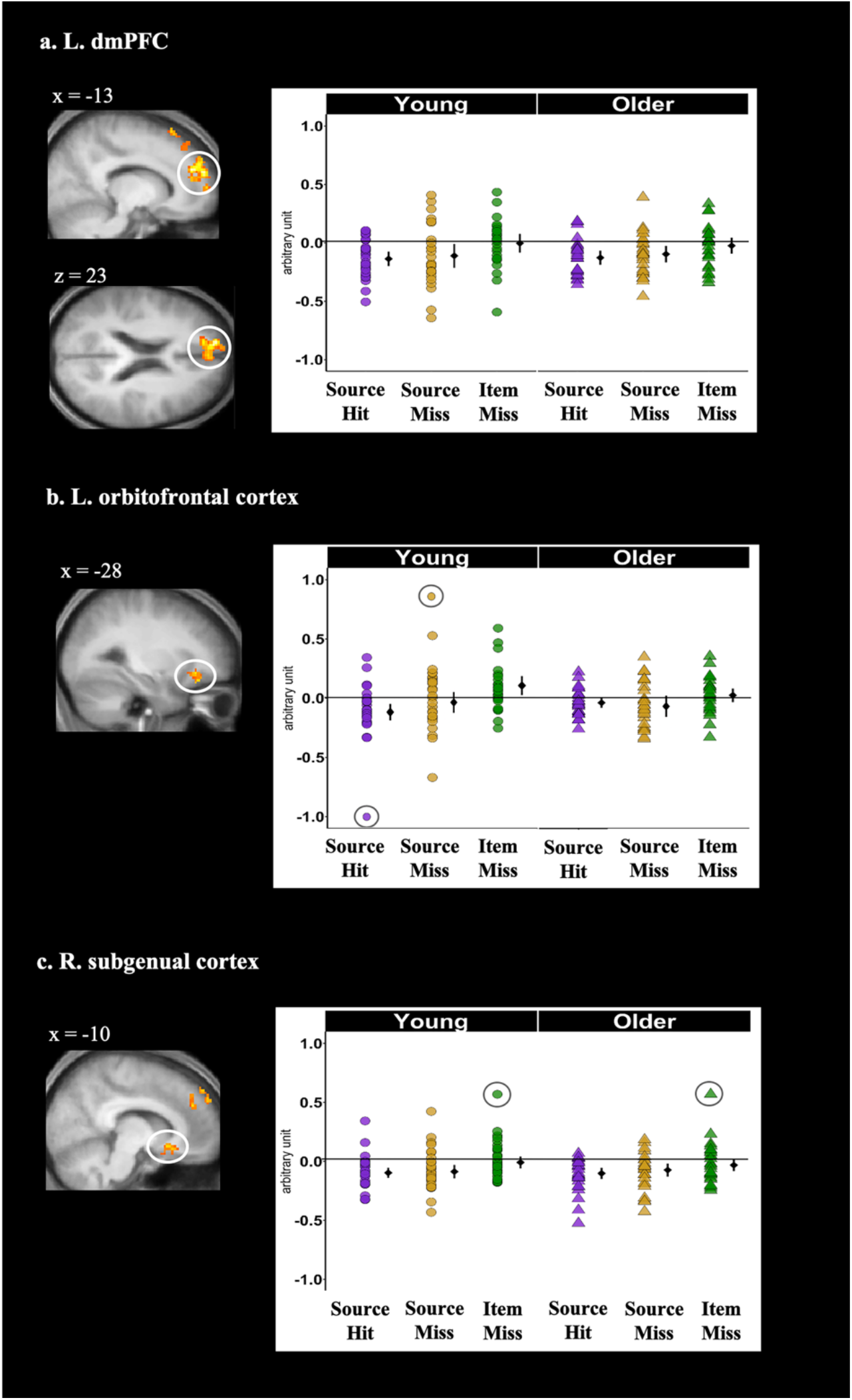
Cortical regions identified by the whole brain analyses that demonstrated significant age-invariant main effects of memory condition during the pre-stimulus period: (a) left dmPFC, (b) left orbitofrontal cortex, and (c) right subgenual cortex. Shown on the left side of each panel are the significant clusters, marked by circles, overlaid on the across-participants mean T1-weighted structural image. Parameter estimates of pre-stimulus activity for each region are plotted on the right side of each panel. Individual parameter estimates (arbitrary units) are plotted separately for young (circles) and older adults (triangles) as a function of subsequent memory. Black dots represent the group means and the error bars signify 95% confidence intervals. The confidence intervals were computed excluding the flagged outliers (list-wise exclusion).

In contrast to the foregoing analyses, age by subsequent memory interactions were observed in both the left angular gyrus and fusiform ROIs (see Figure 3). Follow-up t-tests revealed that, in both regions, pre-stimulus activity was lower for subsequent source hit and source miss trials relative to item misses in young adults (Table 6), whereas no significant differences between memory conditions were identified in older adults.

**Figure 3.**
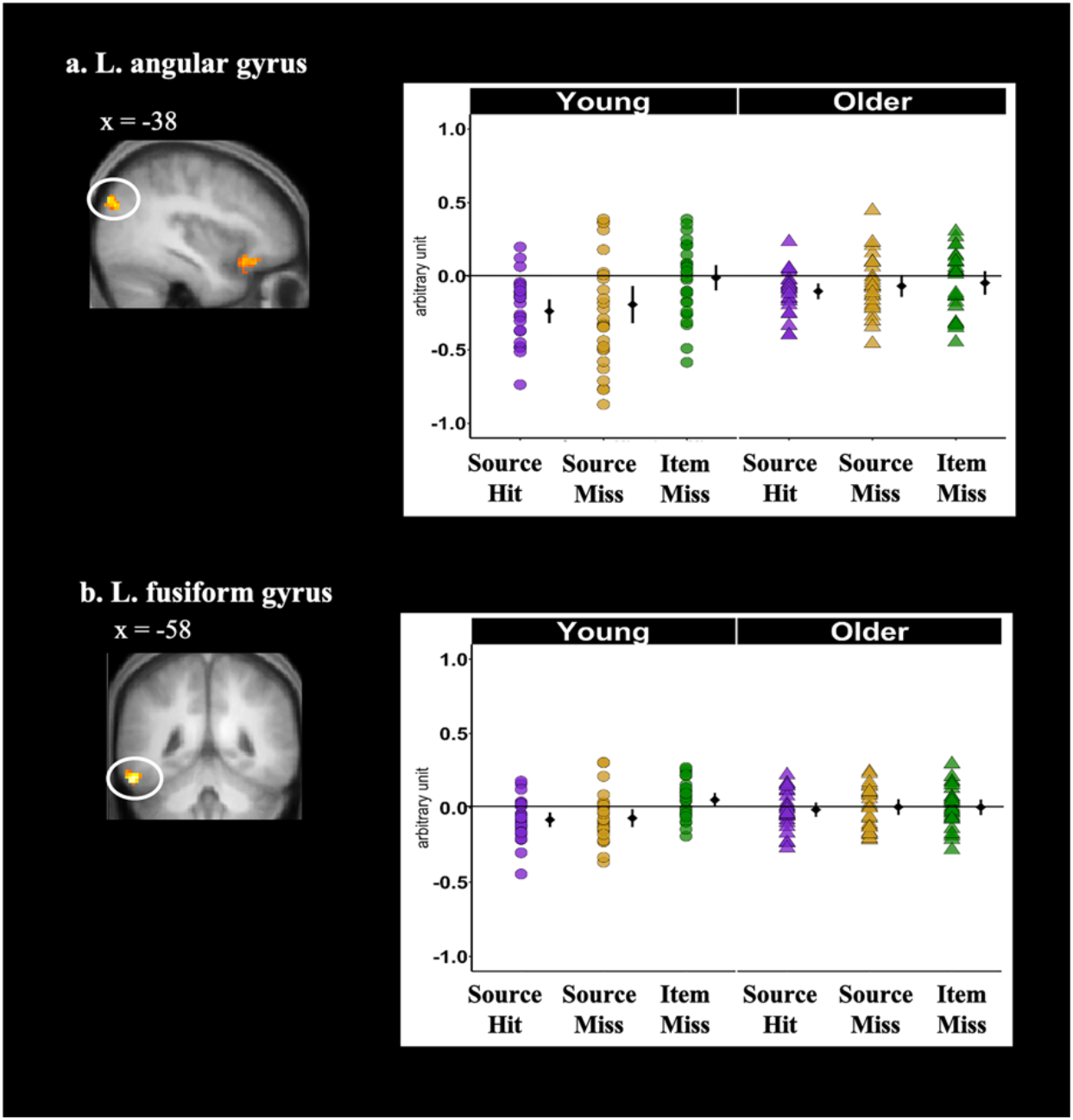
Cortical regions identified by the whole brain analyses that demonstrated significant age-dependent main effects of memory condition during the pre-stimulus period: (a) left angular gyrus and (b) left fusiform gyrus. Shown on the left side of each panel are the significant clusters, marked by circles, overlaid on the across-participants mean T1-weighted structural. Parameter estimates of pre-stimulus activity for each region are plotted on the right side of each panel. Individual parameter estimates (arbitrary units) are plotted separately for young (circles) and older adults (triangles) as a function of subsequent memory. Black dots represent the group means and the error bars signify 95% confidence intervals. The confidence intervals were computed excluding the flagged outliers (list-wise exclusion).

The FIR time-courses (see Methods) are illustrated in Figures 5A and 5B for two representative cortical regions: left dmPFC (the largest of the clusters to demonstrate pre-stimulus SMEs), and left fusiform (one of the two regions where significant age-related differences were identified). As can be seen in the relevant panels of the figure, negative SMEs are evident well before the onset of the study item, indicating that the first level GLMs were successful in identifying pre-stimulus activity.

#### Hippocampus

The small volume corrected second level GLM (see Methods) identified a main effect of subsequent memory in the left anterior hippocampus (−28, −12, −18; peak z = 3.83, 34 voxels; Figure 4). Mean parameter estimates from a 3mm radius sphere centered on the peak of the effect were extracted and subjected to a 2 (age group) by 3 (memory condition) ANOVA. The results revealed a non-significant age group by memory condition interaction (F_1.65, 87.18_ = 0.338, p = 0.714, η_p_^2^ = 0.01), suggesting that pre-stimulus SMEs in the hippocampus were age-invariant. Follow-up contrasts revealed significantly lower activity for source hits relative to item misses (t_54_ = 4.74, p < 0.001; see Figure 3), whereas neither source hits nor item misses differed significantly from source misses. Results did not change after removing the outlying data point (see Figure 3) from the dataset. FIR time-courses illustrating these effects are shown in Figure 5C.

**Figure 4.**
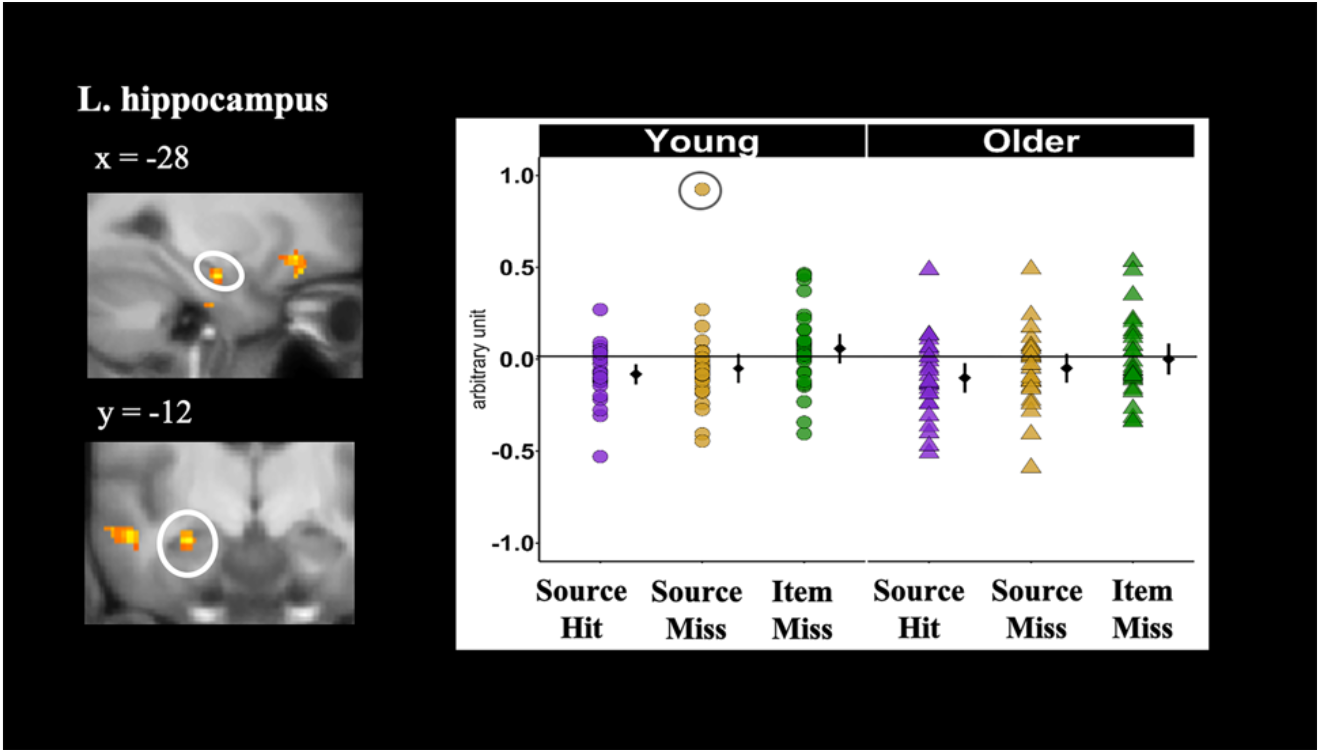
Hippocampal cluster (indicated by the circle) identified by small volume correction that demonstrated a significant pre-stimulus SMEs. The left panel shows the significant cluster overlaid on the across-participants mean T1-weighted structural. The right panel plots the parameter estimates of pre-stimulus activity as a function of age group and subsequent memory. Individual parameter estimates (arbitrary units) are plotted separately for young (circles) and older adults (triangles) as a function of subsequent memory. Black dots represent the group means and the error bars signify 95% confidence intervals. The confidence intervals were computed excluding the flagged outliers (circles).

**Figure 5.**
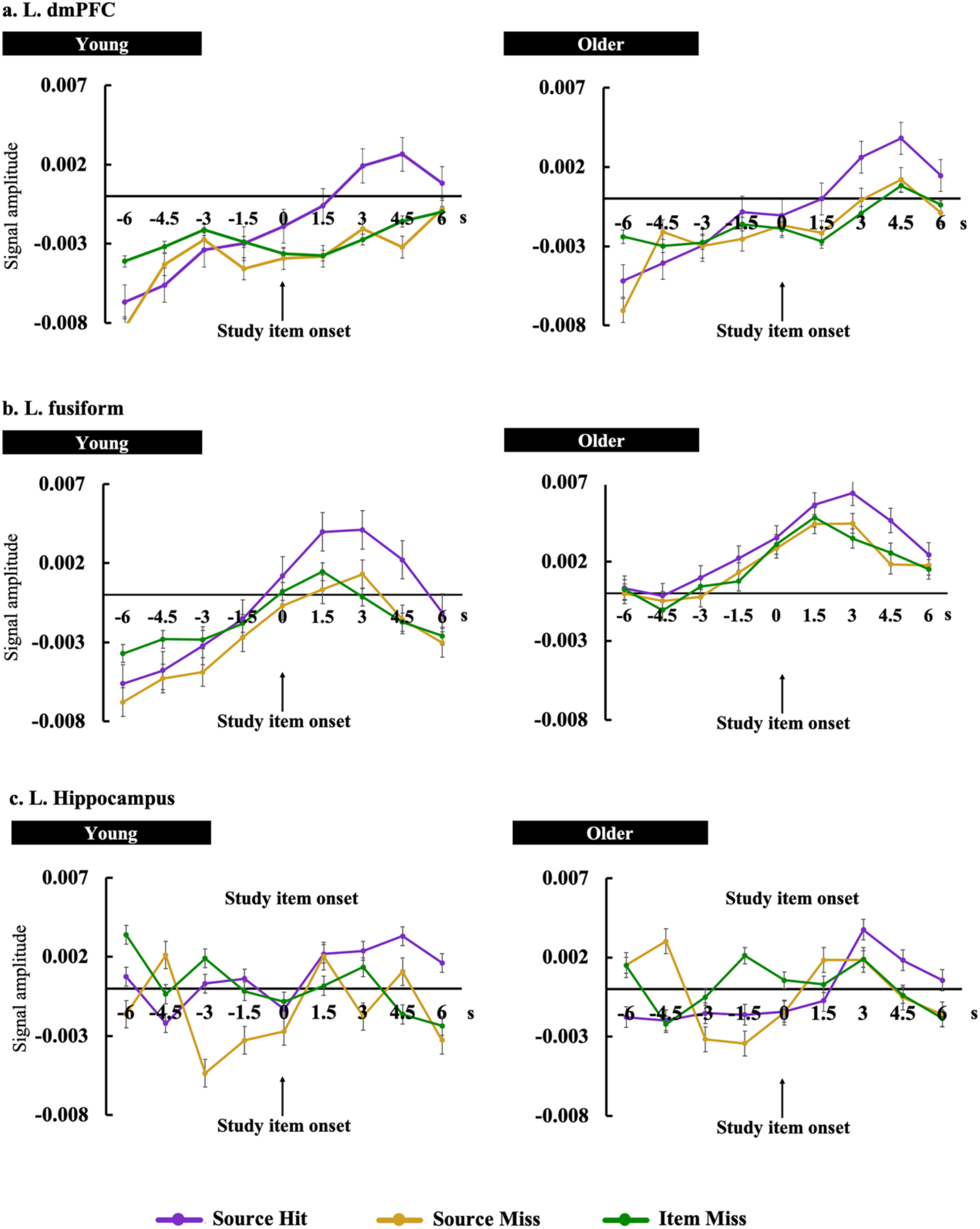
Time-courses of the pre-stimulus subsequent memory effects for young (left panel) and older (right panel) adults in (a) left dmPFC, (b) left fusiform gyrus, and (c) left hippocampus. Signal amplitude is in arbitrary units. Time (in second) is plotted on the x axis, with *time 0*, marked by lines, indicating onset of the study item.

## Discussion

The present experiment investigated pre-stimulus encoding-related neural activity in healthy young and older adults. Age-invariant negative pre-stimulus SMEs were identified in the left hippocampus, dmPFC, orbitofrontal cortex, and right subgenual cortex. In contrast to these age-invariant effects, pre-stimulus SMEs in the left angular gyrus and fusiform cortex were detectable in the young age group only. In general, the effects predicted successful item, rather than source, memory on the subsequent memory test. Below, we discuss the implications of these findings for the understanding of age differences in episodic memory encoding and the functional significance of pre-stimulus SMEs.

### Behavioral Performance

#### Item memory

Collapsed across confidence ratings, neither accuracy nor response bias estimates differed significantly between the age groups. Accuracy for highly confident judgments was also statistically equivalent between the two groups, but older subjects demonstrated a significantly more liberal response bias for these judgments (i.e., older adults were more willing to endorse both old and new items with a high confidence ‘old’ response). Under the assumptions of the two-high threshold model from which the bias estimate was derived (Snodgrass and Corwin, 1988), this finding implies that a higher proportion of the older than the young adults’ confident, correct item judgments were “lucky guesses”. If, instead, a signal detection perspective is adopted (Stretch & Wixted, 1998), the finding implies that the older participants required less mnemonic evidence than the young group before making a high confidence judgment. Either way, the finding indicates that the memory strength of confidently recognized old items was not fully equated between the young and older samples.

#### Source Memory

Consistent with many prior reports (Spencer & Raz, 1995; Old & Naveh-Benjamin, 2008) source memory performance was higher in the young than in the older age group. This age difference was not evident, however, when analysis was restricted to confident source judgments that followed confident item memory judgments (the trial type employed in our fMRI analyses). However, whereas the proportions of confident correct source judgments did not differ according to age (replicating the findings of Mattson et al., 2014), older adults were markedly more prone to make erroneous source judgments with high confidence. Thus, as in the case of item memory judgments, it cannot be assumed that source memory strength was equated between the age groups, even for source judgments made with high confidence.

### fMRI findings

#### Negative pre-stimulus SMEs

In striking contrast to findings from prior fMRI studies employing intentional or semantically elaborative study tasks (e.g., Adcock et al, 2006; Park & Rugg, 2009; Addante et al, 2015; de Chastelaine & Rugg, 2015), here we identified exclusively negative pre-stimulus SMEs in both the cortex and the left hippocampus. The reasons for this striking divergence from prior findings are unclear, although part of the answer may lie in the plethora of procedural differences between the present and prior studies (as noted in the Introduction, a seemingly minor procedural variation can be sufficient to reverse the direction of electrophysiological measures of pre-stimulus SMEs; Koen et al., 2018). For example, to our knowledge, the present study is the first to use fMRI to examine pre-stimulus SMEs predictive of subsequent source, as opposed to item or associative, memory performance. In addition, the present study adopted a slightly different trial structure from that employed in prior studies, in that the informative pre-stimulus cues (signaling which judgment should be made on the upcoming study item) were themselves preceded by a 500 ms duration alerting cue. Thus, unlike in prior studies, here the onset of the informative cues was predictable. Lastly, in the present study the pre-stimulus cues informed participants both about the nature of the upcoming study task and the hand of response. The cue-item interval therefore served as an opportunity for both differential task and differential response preparation. Determining which of these or other procedural differences between the present and prior studies were responsible for the negative pre-stimulus SMEs identified here will require future research.

What might be the functional significance of these SMEs? As was noted in the Introduction, while reported infrequently, negative pre-stimulus SMEs are not unprecedented. Employing a cued study procedure not unlike that used here, de Chastelaine and Rugg (2015) identified positive pre-stimulus SMEs in the medial temporal lobe when an animacy judgment was required on the study words, but negative effects when the task involved a ‘shallow’ syllable judgment. The authors proposed that the negative pre-stimulus SMEs observed in the syllabic study task reflected the detrimental consequences of effective preparation for the upcoming task, which increased the likelihood of encoding information about the study event that was incompatible with the retrieval demands of the subsequent memory test (cf. Morris et al., 1977). [Such “negative transfer” accounts were originally advanced to explain negative cortical and hippocampal post-stimulus SMEs (e.g., Otten and Rugg, 2001; Shrager et al., 2008; Staresina & Davachi, 2008; Hill et al., 2020]. From this perspective, therefore, the present negative pre-stimulus SMEs reflect the deleterious consequences of the successful encoding of study information incompatible with, or inaccessible to, the subsequent memory test. It is not obvious, however, why this would be the case here given that prior studies employing similar semantically elaborative study tasks found no evidence of negative pre-stimulus SMEs.

An alternative account follows from the findings of another study in which negative pre-stimulus SMEs were reported. As was described in the Introduction, in an un-cued paradigm, Yoo et al. (2012) reported that study items (scenes) were more likely to be remembered when the fMRI BOLD signal preceding the presentation of the items was of relatively low amplitude (for related EEG findings, see Salari & Rose, 2016). The authors proposed that memory for a study event benefits when regions necessary for its processing are relatively quiescent at the time of its occurrence, allowing the neural resources of the regions to be allocated more fully to the event. Adapting this account to the present findings, it could be argued that the negative pre-stimulus SMEs that were observed here are a reflection of the same kinds of spontaneous fluctuations in ‘neural state’ that underpinned Yoo et al.’s findings (see also Otten et al., 2002), or alternately that the beneficial ‘down-regulation’ of pre-stimulus activity was an active process triggered by the presentation of the pre-stimulus cues. While both of these accounts are arguably attractive, either version faces the same challenge mentioned in the prior paragraph, namely, the failure of analogous prior studies to identify negative pre-stimulus SMEs.

#### Effects of age on pre-stimulus SMEs

Contrary to our pre-experimental prediction, robust but age-invariant pre-stimulus SMEs were identified in the left hippocampus as well as in three of the five cortical regions where pre-stimulus SMEs were observed. Although it has been reported that older adults demonstrate reduced pre-stimulus SMEs (Koen et al., 2018), findings of age-invariant pre-stimulus SMEs are not unprecedented. Notably, Strunk & Duarte (2019) reported age-invariant pre-stimulus SMEs in oscillatory neural activity elicited by informative cues that signaled the modality of the upcoming study item. The authors interpreted these findings as evidence for intact pre-stimulus preparatory processes in their older adults.

The significance of the present age-invariant findings depends heavily on the interpretation given to the pre-stimulus SMEs that were identified (see above). If the effects are interpreted as reflections of active preparation for the upcoming study event, then they constrain the proposal of Koen et al (2018) that older adults are less able (or less willing) to engage proactive processing in anticipation of an upcoming event: in some neural regions at least, including those known to play a necessary role in episodic encoding, such as the hippocampus, older and young adults appear to engage such processing to an equivalent extent. Alternately, if the present effects are interpreted in terms of spontaneous, involuntary fluctuations in neural state (cf. Yoo et al., 2012), then they do not speak to this issue. Regardless of which interpretation is correct, however, it is clear that, as is the case for positive post-stimulus SMEs (see Introduction), negative pre-stimulus SMEs can be remarkably resilient to the effects of age.

In contrast to the above-mentioned regions, pre-stimulus effects in two other cortical regions – left fusiform cortex and angular gyrus – were detectable in young adults only. These findings are consistent both with our prediction that we would identify age-related attenuation of pre-stimulus SMEs (see Introduction) and with the findings from Koen et al. (2018). However, while the present findings might signify a failure on the part of older adults to engage preparatory or proactive processes in these regions, they do not in themselves support the proposal that older adults cannot engage such processes to the same extent as their young counterparts: as was discussed in the preceding paragraph, pre-stimulus effects in other cortical regions and the hippocampus were equally evident in the two age groups. From the standpoint of a ‘preparatory’ interpretation, this regional heterogeneity in the pattern of preserved and attenuated pre-stimulus SMEs in older adults suggests that a generic construct such as reduced task preparation is ill-suited to accounting for age differences in pre-stimulus encoding-related activity.

#### Patterning of pre-stimulus SMEs

Prior fMRI studies of pre-stimulus SMEs employed either Remember/Know/New (Adcock et al, 2006; Park & Rugg, 2009; de Chastelaine & Rugg, 2015) or associative recognition procedures (Addante et al, 2015) to examine subsequent memory performance, and invariably reported pre-stimulus SMEs that were selectively predictive of recollection-rather than familiarity-based memory judgments. In the present study, other than in the orbitofrontal cortex and hippocampus, pre-stimulus SMEs effects took the form of reduced activity for both source hits *and* source misses relative to item misses; that is, pre-stimulus SMEs in these regions seemingly predicted item recognition rather than source recollection. To the extent that source misses were recognized largely on the basis of familiarity, the present findings therefore run counter to those reported in prior studies. However, although the present findings might indeed indicate that pre-stimulus processes can enhance encoding that supports familiarity-based memory judgments, this is not the only possible interpretation. The source miss bin in this study comprised items that attracted high confidence item hits, and it has been demonstrated that high confident item responses are usually associated with subjective reports of recollection (Yonelinas, 2001; Koen & Yonelinas, 2010). Hence, in the current study it is highly likely that a significant proportion of items classified as source misses were recognized not only on the basis their familiarity, but also because they elicited a ‘non-criterial’ recollection signal (i.e., recollection of information about the study event that was non-diagnostic of source; cf. Yonelinas & Jacoby, 1996; Parks, 2007). Furthermore, the source miss category included items attracting correct, low confidence source judgments (see fMRI analysis). Although the accuracy of these judgments was markedly lower than that of the judgments made with high confidence, it was not at chance levels (see Table 4). Thus, it is possible that the tendency in most regions for pre-stimulus SMEs to predict successful item memory judgments on the later memory test is a reflection of the fact that, for a substantial proportion of the study trials categorized as source misses, episodic encoding was not entirely unsuccessful. By this account, the graded pre-stimulus SMEs identified in the hippocampus and orbitofrontal cortex suggest that pre-stimulus activity in these regions is more predictive of the amount of episodic information encoded during the upcoming study event than is the activity in other regions where pre-stimulus SMEs were identified (cf. Rugg et al., 2012).

#### Limitations

The present study has several limitations. First, it employed a cross-sectional, extreme age-group design. Thus, it not possible to determine whether the differences between the two age groups are a consequence of aging as opposed to one or more factors that are confounded with age, such as cohort effects (Rugg, 2017). Second, the rationale for the use of confidence ratings for the item and source memory judgments was to allow examination of age differences in pre-stimulus SMEs in the absence of the confounding factor of age-related differences in memory strength (see Rugg, 2017 for discussion of this and related factors). As was discussed above, we were not entirely successful in this aim, and thus we cannot rule out the possibility that the age differences identified in the angular gyrus and fusiform cortex were influenced by this factor, if only partially. Lastly, regionally specific age differences in the transfer function mediating between neural activity and the fMRI BOLD response have been reported (e.g. Lu et al., 2011; Liu et al., 2013). Thus, age differences in neurovascular coupling are a potentially confounding factor in the present study. Whereas these differences are arguably of relatively little concern when considering null effects of age of the present pre-stimulus SMEs, it is possible that they contributed to the age differences identified in the two posterior cortical regions noted above.

#### Conclusions

The present findings extend prior reports of pre-stimulus SMEs and provide initial insights into age-related differences in these effects as they are expressed in fMRI BOLD signals. The finding of regional differences in the impact of age on pre-stimulus SMEs, with some regions demonstrating age-invariant effects, and other effects that were exclusive to young adults, suggests that, as is the case for post-stimulus SMEs, pre-stimulus SMEs are unlikely to be explained by appeal to a single, general construct such as engagement of preparatory task sets (cf. Koen et al., 2018).

## Acknowledgments

We are grateful to Christopher Hawkins for his assistance with data collection and our colleagues at the University of Texas Southwestern Medical Center Advanced Imaging Research Center for their assistance with fMRI data acquisition.

## References

Adcock, R. A., Thangavel, A., Whitfield-Gabrieli, S., Knutson, B., & Gabrieli, J. D. E. (2006). Reward-Motivated Learning: Mesolimbic Activation Precedes Memory Formation. Neuron, 50(3), 507–517.

Addante, R. J., de Chastelaine, M., & Rugg, M. D. (2015). Pre-stimulus neural activity predicts successful encoding of inter-item associations. NeuroImage, 105, 21–31.

Benton, A. L. (1968). Differential behavioral effects in frontal lobe disease. Neuropsychologia, 6(1), 53–60.

Braver, T. S. (2012). The variable nature of cognitive control: a dual mechanisms framework. Trends in Cognitive Sciences, 16(2), 106–113.

Braver, T. S., Paxton, J. L., Locke, H. S., & Barch, D. M. (2009). Flexible neural mechanisms of cognitive control within human prefrontal cortex. Proceedings of the National Academy of Sciences of the United States of America, 106(18), 7351–7356.

Brewer, J. B., Zhao, Z., Desmond, J. E., Glover, G. H., & Gabrieli, J. D. E. (1998). Making memories: Brain activity that predicts how well visual experience will be remembered. Science, 281(5380), 1185–1187.

Chumbley, J., Worsley, K., Flandin, G., & Friston, K. (2010). Topological FDR for neuroimaging. NeuroImage, 49(4), 3057–3064.

Cohen, N., Ben-Yakov, A., Weber, J., Edelson, M. G., Paz, R., & Dudai, Y. (2019). Prestimulus Activity in the Cingulo-Opercular Network Predicts Memory for Naturalistic Episodic Experience. Cerebral Cortex, 1–12.

Cohen, N., Pell, L., Edelson, M. G., Ben-Yakov, A., Pine, A., & Dudai, Y. (2015). Peri-encoding predictors of memory encoding and consolidation. Neuroscience and Biobehavioral Reviews, 50, 128–142.

Craik, F. I. M., & Rose, N. S. (2012). Memory encoding and aging: A neurocognitive perspective. Neuroscience and Biobehavioral Reviews, 36(7), 1729–1739.

Craik, F. I., & Byrd, M. (1982). Aging and cognitive deficits. In Aging and cognitive processes (pp.191–211). Springer, Boston, MA.

de Chastelaine, M., Mattson, J. T., Wang, T. H., Donley, B. E., & Rugg, M. D. (2016). The relationships between age, associative memory performance, and the neural correlates of successful associative memory encoding. Neurobiology of Aging, 42, 163–176.

de Chastelaine, M., & Rugg, M. D. (2015). The effects of study task on prestimulus subsequent memory effects in the hippocampus. Hippocampus, 25(11), 1217–1223.

de Chastelaine, M., Wang, T. H., Minton, B., Muftuler, L. T., & Rugg, M. D. (2011). The Effects of Age, Memory Performance, and Callosal Integrity on the Neural Correlates of Successful Associative Encoding. Cerebral Cortex, 21(9), 2166–2176.

Dennis, N.A., Hayes, S.M., Prince, S.E., Madden, D.J., Huettel, S.A., & Cabeza, R. (2008). Effects of aging on the neural correlates of successful item and source memory encoding. Journal of experimental psychology: Learning, Memory, and Cognition, 34, 791–808.

Delis, D. C., Kramer, J. H., Kaplan, E., & Ober, B. A. (2000). California verbal learning test (2nd ed.). San Antonio, TX: Psychological Corporation.

Ezzyat, Y., Kragel, J. E., Burke, J. F., Levy, D. F., Lyalenko, A., Wanda, P.,…Kahana, M. J. (2017). Direct Brain Stimulation Modulates Encoding States and Memory Performance in Humans. Current Biology, 27(9), 1251–1258.

Fernández, G., Brewer, J. B., Zhao, Z., Glover, G. H., & Gabrieli, J. D. E. (1999). Level of sustained entorhinal activity at study correlates with subsequent cued-recall performance: A functional magnetic resonance imaging study with high acquisition rate. Hippocampus, 9(1), 35–44.

Friedman, D., & Johnson, R. (2014). Inefficient Encoding as an Explanation for Age-Related Deficits in Recollection-Based Processing. Journal of Psychophysiology, 28(3), 148–161.

Gutchess, A., Welsh, R.C., Hedden, T., Bangert, A., Minear, M., Liu, L., & Park, D. (2005). Aging and the neural correlates of successful picture encoding: frontal activations compensate for decreased medial-temporal activity. Journal of Cognitive Neuroscience, 17, 84–96.

Hill, P. F., King, D. R., Lega, B. C., & Rugg, M. D. (2020). Comparison of fMRI correlates of successful episodic memory encoding in temporal lobe epilepsy patients and healthy controls. NeuroImage, 207(May 2019), 116397.

Koen, J. D., Horne, E. D., Hauck, N., & Rugg, M. D. (2018). Age-related Differences in Prestimulus Subsequent Memory Effects Assessed with Event-related Potentials. Journal of Cognitive Neuroscience, 30(6), 829–850.

Koen, J. D., & Yonelinas, A. P. (2010). Memory variability is due to the contribution of recollection and familiarity, not to encoding variability. Journal of Experimental Psychology: Learning, Memory, and Cognition, 36(6), 1536.

Koen, J. D., & Yonelinas, A. P. (2014). The Effects of Healthy Aging, Amnestic Mild Cognitive Impairment, and Alzheimer’s Disease on Recollection and Familiarity: A Meta-Analytic Review. Neuropsychology Review, 24(3), 332–354.

Liu, P., Hebrank, A. C., Rodrigue, K. M., Kennedy, K. M., Section, J., Park, D. C., & Lu, H. (2013). Age-related differences in memory-encoding fMRI responses after accounting for decline in vascular reactivity. NeuroImage, 78, 415–425.

Lu, H., Xu, F., Rodrigue, K. M., Kennedy, K. M., Cheng, Y., Flicker, B.,…Park, D. C. (2011). Alterations in Cerebral Metabolic Rate and Blood Supply across the Adult Lifespan. Cerebral Cortex, 21(6), 1426–1434.

Maillet, D., & Rajah, M. N. (2014). Dissociable roles of default-mode regions during episodic encoding. NeuroImage, 89, 244–255.

Mattson, J. T., Wang, T. H., De Chastelaine, M., & Rugg, M. D. (2014). Effects of age on negative subsequent memory effects associated with the encoding of item and item-context information. Cerebral Cortex, 24(12), 3322–3333.

Miller, S.L., Celone, K., DePeau, K., Diamond, E., Dickerson, B.C., Rentz, D., Pihlajamäki, M., & Sperling, R.A. (2008). Age-related memory impairment associated with loss of parietal deactivation but preserved hippocampal activation. Proc. Natl. Acad. Sci. USA, 105, 2181–2186.

Morcom, A.M., Good, C.D., Frackowiak, R.S., & Rugg, M.D. (2003). Age effects on the neural correlates of successful memory encoding. Brain, 126, 213–229.

Mormino, E.C., Brandel, M.G., Madison, C.M., Marks, S., Baker, S.L., & Jagust, W.J. (2012). Ab deposition in aging is associated with increases in brain activation during successful memory encoding. Cerebral Cortex, 22, 1813–1823.

Morris, C.D., Bransford, J.D., & Franks, J.J. (1977). Levels of processing versus transfer appropriate processing. Journal of Verbal Learning and Verbal Behavior, 16, 519–533.

Nilsson, L. G. (2003). Memory function in normal aging. Acta Neurologica Scandinavica, Supplement, 107(179), 7–13.

Nyberg, L., Lövdén, M., Riklund, K., Lindenberger, U., & Bäckman, L. (2012). Memory aging and brain maintenance. Trends in Cognitive Sciences, 16(5), 292–305.

Old, S. R., & Naveh-Benjamin, M. (2008). Differential Effects of Age on Item and Associative Measures of Memory: A Meta-Analysis. Psychology and Aging.

Otten, L. J., Henson, R. N. A., & Rugg, M. D. (2002). State-related and item-related neural correlates of successful memory encoding. Nature Neuroscience, 5(12), 1339–1344.

Otten, L. J., Quayle, A. H., Akram, S., Ditewig, T. A., & Rugg, M. D. (2006). Brain activity before an event predicts later recollection. Nature Neuroscience, 9(4), 489–491.

Otten, L. J., Quayle, A. H., & Puvaneswaran, B. (2010). Prestimulus subsequent memory effects for auditory and visual events. Journal of Cognitive Neuroscience, 22(6), 1212–1223.

Otten, L. J., & Rugg, M. D. (2001). When more means less: Neural activity related to unsuccessful memory encoding. Current Biology. https://doi.org/10.1016/S0960-9822(01)00454-7

Padovani, T., Koenig, T., Brandeis, D., & Perrig, W. J. (2011). Different brain activities predict retrieval success during emotional and semantic encoding. Journal of Cognitive Neuroscience, 23(12), 4008–4021.

Paller, K. A., & Wagner, A. D. (2002). Observing the transformation of experience into memory. Trends in Cognitive Sciences, 6(2), 93–102.

Park, H., Kennedy, K.M., Rodrigue, K.M., Hebrank, A., & Park, D.C. (2013). An fMRI study of episodic encoding across the lifespan: changes in subsequent memory effects are evident by middle-age. Neuropsychologia, 51, 448–456.

Park, D. C., Polk, T. A., Mikels, J. A., Taylor, S. F., & Marshuetz, C. (2001). Cerebral aging: integration of brain and behavioral models of cognitive function. Dialogues in clinical neuroscience, 3(3), 151–165.

Park, H., & Rugg, M. D. (2009). Prestimulus hippocampal activity predicts later recollection. Hippocampus, 20(1), 24–28.

Parks, C. M. (2007). The role of noncriterial recollection in estimating recollection and familiarity. Journal of Memory and Language, 57(1), 81–100.

Raven J., Raven J. C., Courth J. H. (2000). Manual for Raven’s progressive matrices and vocabulary scales. Section 4: the advanced progressive matrices. San Antonio, TX: Harcourt Assessment

Reitan, R. M., & Wolfson, D. (1985). The Halstead–Reitan neuropsychological test battery: Therapy and clinical interpretation. Tucson, AZ: Neuropsychological Press.

Rugg, M. D. (2017). Interpreting age-related differences in memory-related neural activity. In R. Cabeza, L. Nyberg, & D. C. Park (Eds.), Cognitive Neuroscience of Aging: Linking Cognitive and Cerebral Aging (2nd ed., pp. 183–206). Oxford University Press.

Rugg, M. D., Vilberg, K. L., Mattson, J. T., Yu, S. S., Johnson, J. D., & Suzuki, M. (2012). Item memory, context memory and the hippocampus: FMRI evidence. Neuropsychologia, 50(13), 3070–3079.

Sadeh, T., Chen, J., Goshen-Gottstein, Y., & Moscovitch, M. (2019). Overlap between hippocampal pre-encoding and encoding patterns supports episodic memory. Hippocampus, 29(9), 836–847.

Salari, N., & Rose, M. (2016). Dissociation of the functional relevance of different pre-stimulus oscillatory activity for memory formation. Neuroimage, 125, 1013–1021.

Schneider, S. L., & Rose, M. (2016). Intention to encode boosts memory-related pre-stimulus EEG beta power. Neuroimage, 125, 978–987.

Shrager, Y., Kirwan, C.B., Squire, L.R., 2008. Activity in both hippocampus and perirhinal cortex predicts the memory strength of subsequently remembered information. Neuron, 59 (4), 547–553.

Smith, A. (1973). Symbol digit modalities test. Los Angeles: Western Psychological Services.

Snodgrass, J. G., & Corwin, J. (1988). Pragmatics of Measuring Recognition Memory:Applications to Dementia and Amnesia. Journal of Experimental Psychology: General, 117(1), 34–50.

Spencer, W. D., & Raz, N. (1995). Differential effects of aging on memory for content and context: a meta-analysis. Psychology and Aging, 10(4), 527–539.

Spreen, O., & Benton, A. L. (1977). Neurosensory center comprehensive examination for aphasia. Victoria, Canada: Neuropsychology Laboratory.

Squire, L. R., Wixted, J. T., & Clark, R. E. (2007). Recognition memory and the medial temporal lobe: A new perspective. Nature Reviews Neuroscience. 8(11), 872–883.

Staresina, B.P., & Davachi, L. (2008). Selective and shared contributions of the hippocampus and perirhinal cortex to episodic item and associative encoding. Journal of Cognitive Neuroscience, 20, 1478–1489.

Stretch, V., & Wixted, J. T. (1998). Decision rules for recognition memory confidence judgments. Journal of Experimental Psychology: Learning Memory and Cognition.

Strunk, J., & Duarte, A. (2019). Prestimulus and poststimulus oscillatory activity predicts successful episodic encoding for both young and older adults. Neurobiology of Aging, 77, 1–12.

Sweeney-Reed, C. M., Zaehle, T., Voges, J., Schmitt, F. C., Buentjen, L., Kopitzki, K.,… & Rugg, M. D. (2016). Pre-stimulus thalamic theta power predicts human memory formation. Neuroimage, 138, 100–108.

Wagner, A. D. (1998). Building Memories: Remembering and Forgetting of Verbal Experiences as Predicted by Brain Activity. Science, 281(5380), 1188–1191.

Wechsler, D. (1981). WAIS-R: Wechsler Adult Intelligence Scale-Revised. New York: Psychological Corporation.

Wechsler, D. (2009). Wechsler Memory Scale (4th ed.). San Antonio, TX: Psychological Corporation.

Werkle-Bergner, M., Müller, V., Li, S.-C., & Lindenberger, U. (2006). Cortical EEG correlates of successful memory encoding: Implications for lifespan comparisons. Neuroscience & Biobehavioral Reviews, 30(6), 839–854.

Yonelinas, A. P., & Jacoby, L. L. (1996). Noncriterial Recollection: Familiarity as Automatic, Irrelevant Recollection. Consciousness and Cognition, 5(1-2), 131–141.

Yonelinas, A. P. (2001). Consciousness, control, and confidence: the 3 Cs of recognition memory. Journal of Experimental Psychology: General, 130(3), 361.

Yoo, J. J., Hinds, O., Ofen, N., Thompson, T. W., Whitfield-Gabrieli, S., Triantafyllou, C., & Gabrieli, J. D. E. (2012). When the brain is prepared to learn: Enhancing human learning using real-time fMRI. Neuroimage, 59(1), 846–852.

